# Autoregulation of pre-mRNA processing for buffering noisy gene expression

**DOI:** 10.1101/623181

**Authors:** Madeline Smith, Khem Raj Ghusinga, Abhyudai Singh

## Abstract

Stochastic variation in the level of a protein among cells of the same population is ubiquitous across cell types and organisms. These random variations are a consequence of low-copy numbers, amplified by the characteristically probabilistic nature of biochemical reactions associated with gene-expression. We systematically compare and contrast negative feedback architectures in their ability to regulate random fluctuations in protein levels. Our stochastic model consists of gene synthesizing pre-mRNAs in transcriptional bursts. Each pre-mRNA transcript is exported to the cytoplasm and is subsequently translated into protein molecules. In this setup, three feedbacks architectures are implemented: protein inhibiting transcription of its own gene (I), protein enhancing the nuclear pre-mRNA decay rate (II), and protein inhibiting the export of pre-mRNAs (III). Explicit analytic expressions are developed to quantify the protein noise levels for each feedback strategy. Mathematically controlled comparisons provide insights into the noise-suppression properties of these feedbacks. For example, when the protein half-life is long, or the pre-mRNA decay is fast, then feedback architecture I provides the best noise attenuation. In contrast, when the timescales of export, mRNA, and protein turnover are similar, then III is superior to both II and I. We finally discuss biological relevance of these findings in context of noise suppression in an HIV cell-fate decision circuit.

## I. INTRODUCTION

Stochastic variation in the level of protein among cells of the same population is ubiquitous across cell types and organisms. These random variations are a consequence of low-copy numbers, amplified by the characteristically probabilistic nature of biochemical reactions associated with gene-expression [1], [2]. This variation is often referred to as gene-expression noise. The prevalence of this noise suggests that cells should have mechanisms to cope with it, suppress it, or even utilize it to their advantage [3]–[15].

There is immense interest in how cells suppress gene-expression noise. Previous studies investigate regulatory mechanisms that cells may employ by incorporating feedback into models of gene-expression [16]–[29]. These models typically include the gene promoter, mRNA, and protein, and are thus limited to transcriptional or translational feedback. However, recent experimental studies have found that both un-spliced, precursor mRNA as well as stable, cytoplasmic mRNA play a role in the regulation of gene-expression [30]–[38]. This insight challenges simpler models and suggests the possibility of feedback strategies that include the precursor mRNA species.

In this work, we investigate negative feedback mechanisms in their ability to suppress noise in a model of gene-expression that distinguishes precursor mRNA from cytoplasmic mRNA. Briefly, our model assumes that each gene synthesizes pre-mRNAs in the nucleus in transcriptional bursts. Modeling transcription by bursting allows incorporation of the active (ON) and inactive (OFF) intervals of promoter activity [39]–[46]. Then, each pre-mRNA transcript is exported from the nucleus to the cytoplasm. The cytoplasmic mRNA is subsequently translated into protein molecules. This framework allows us to explore additional regulatory strategies involving the precursor mRNA.

We specifically consider the following three feedbacks:

I. the protein inhibits transcription of its own gene,
II. the protein enhances the nuclear pre-mRNA degradation rate, and
III. the protein inhibits the export of pre-mRNAs.

We compare and contrast the noise suppression abilities of these three forms of negative feedback in a mathematically controlled fashion [47]. To that end, the mean level of protein is kept constant among all architectures. Our analysis reveals when pre-mRNA export occurs at a higher rate than its degradation, feedback architecture I provides the best noise attenuation. In contrast, when mRNA is unstable or when pre-mRNA is more likely to degrade than export, then III is superior to both II and I for high feedback strength.

The paper is organized as follows. Section II introduces a gene-expression model without feedback regulation and and uses differential equations for its moments to quantify noise in protein level. In Section III, we modify the model, incorporate the aforementioned three feedback strategies, and compute the noise in protein level. Next, in Section IV, we compare and contrast noise suppression of feedback architectures I-III under different regimes. A discussion of biological relevance of these findings in context of the role noise suppression plays in HIV concludes the paper in Section V.

## II. GENE-EXPRESSION MODEL WITH NO REGULATION

We model gene-expression starting with nuclear pre-mRNA transcripts produced in bursts. Transcriptional bursting at the gene promoter occurs at a rate of *k*_*r*_, creating *B* number of pre-mRNA transcripts, where *B* is a discrete, positive-valued, random variable with distribution

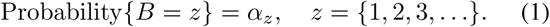

The pre-mRNA is spliced and exported out of the nucleus at a rate of *k*_*m*_ to become functional mRNA. It is then translated into protein at a rate *k*_*p*_. The model additionally includes degradation rates of pre-mRNA (*γ*_*r*_), mRNA (*γ*_*m*_), and protein (*γ*_*p*_).

Let *r*(*t*), *m*(*t*), and *p*(*t*) represent the count of pre-mRNA, mRNA, and protein molecules at time *t*, accordingly. The probabilities of occurrences of these events in an infinitesimal time-interval (*t, t* + *dt*) are described in Table 1.

**TABLE I:** Gene-expression events and corresponding rates of occurrence.

| Event | Change in Population Count | Probability Event will Occur in $(t, t + dt)$ |
| --- | --- | --- |
| Transcriptional Bursting | $r(t) \rightarrow r(t) + B$ | $k_r dt$ |
| pre-mRNA Degradation | $r(t) \rightarrow r(t) - 1$ | $\gamma_r r(t) dt$ |
| pre-mRNA Export | $r(t) \rightarrow r(t) - 1$<br>$m(t) \rightarrow m(t) + 1$ | $k_m r(t) dt$ |
| mRNA Degradation | $m(t) \rightarrow m(t) - 1$ | $\gamma_m m(t) dt$ |
| Protein Translation | $p(t) \rightarrow p(t) + 1$ | $k_p m(t) dt$ |
| Protein Degradation | $p(t) \rightarrow p(t) - 1$ | $\gamma_p p(t) dt$ |

To quantify the noise in protein levels of the gene-expression model, we write differential equations for statistical moments. For the gene-expression model without feedback, the time derivative of the expected value of a differentiable function *φ*(*r, m, p*) is given by [48], [49]

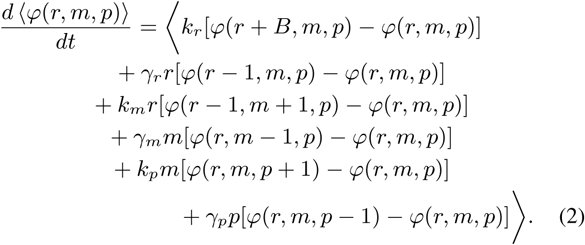

Here, and for the remainder of the paper, ⟨.⟩ is used to represent the expected value.

To compute moment dynamics, *φ* is chosen as a corresponding monomial. We are particularly interested in studying the noise in protein level, so we find the protein steadystate moments by equating its first and second moment (corresponding to the mean and variance in protein level) to zero. This allows for quantification of protein level noise via the coefficient of variation squared (*CV* ^2^, variance divided by squared of the mean) [50].

We introduce the following four parameters for simplifying the moments’ expressions

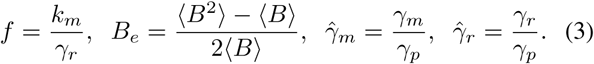

Here, *f* is the ratio of pre-mRNA export rate from the nucleus to its degradation rate; *B*_*e*_ is related to transcriptional burst size; 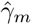 is the mRNA degradation rate normalized by the protein degradation rate; and 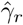 is the pre-mRNA degradation rate normalized by the protein degradation rate.

Solving for steady-state moments, and substituting the above parameters yields

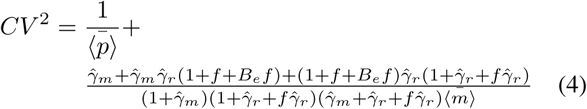

Where 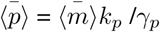 is the steady-state mean protein level. Note that in (4), the first term is the intrinsic Poissonian noise due to stochastic births and deaths of single protein molecules. This term can be ignored when considering the order of magnitude size difference in population counts of mRNA and protein 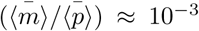 [2]. The resulting simplified coefficient of variation squared is depicted in the center column of Fig. 2.

**Fig. 1:**
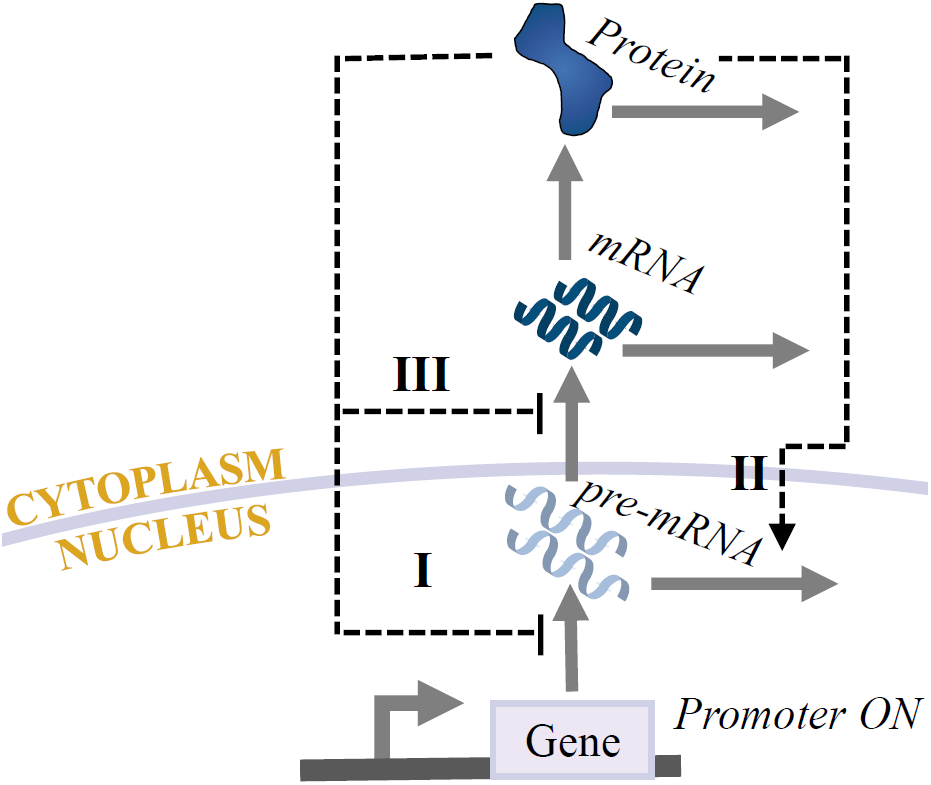
Process of gene-expression modeled by precursor mRNA produced in transcriptional bursts inside the nucleus from the active gene promoter. Precursor mRNA is spliced and exported out of the nucleus as functional mRNA, and finally translated into protein. Proposed feedback mechanisms affecting pre-mRNA transcription (I), degradation (II), or export (III) are depicted as dashed lines. The arrow head denotes activation, while the flat arrow denotes inhibition.

**Fig. 2:**
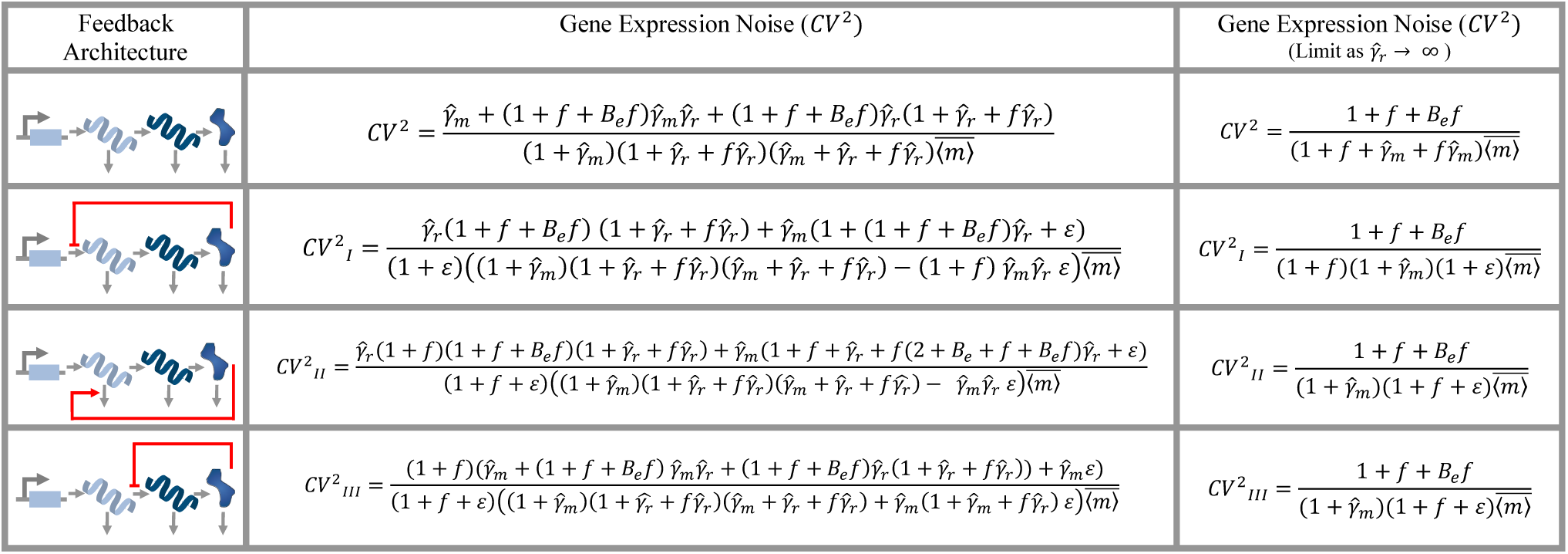
Gene-expression noise of each feedback architecture is quantified by the coefficient of variation squared 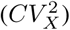, located in the center column of the table. The leftmost column shows the form of feedback present: no feedback, (I) protein inhibition of its own transcription, (II) protein enhancing nuclear pre-mRNA degradation, and (III) protein inhibition of pre-mRNA export. The rightmost column contains 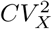 for a limiting case of fast pre-mRNA degradation.

We next analyze gene-expression noise in different limits. First, we consider limiting cases of *f*. When *f* ≫ 1, premRNA is more likely to be exported from the nucleus, and the resulting noise equation simplifies to

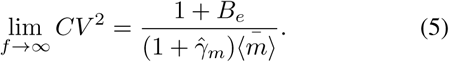

In the opposite case of *f* ≪ 1, the noise equation becomes

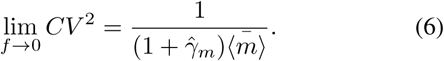

Interestingly, in this case the noise becomes independent of the burst size as opposed to the *f* ≫ 1 case, wherein the noise depends upon the burst size. In both cases, however, the noise in protein level depends inversely on the average steady-state level of mRNA 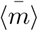.

## III. INTRODUCING REGULATORY MECHANISMS IN GENE-EXPRESSION

Here, we implement three different feedback architectures and systematically compare their abilities to suppress noise. We specifically consider three mechanisms: the protein inhibits transcription of its own gene (I), the protein enhances the nuclear pre-mRNA degradation rate (II), and the protein inhibits the export of pre-mRNAs (III). The architecture I is well-known as negative autoregulation, e.g., see [51]. We consider feedbacks II and III which involve pre-mRNAs and therefore are specific to our model formulation. The feedback architecture II is motivated from regulation of the PABPN1 gene which has been suggested to reduce the spliced vs. unspliced ratio of pre-mRNAs, thus reducing PABPN1’s own gene-expression and ultimately decreasing noise [52]. The last form of feedback (III) has also been suggested in the NUP153 gene whose over-expression slows down the transcript export from the nucleus, resulting in lower variability while maintaining the original steady-state protein count [36], [37], [53].

We incorporate feedback by no longer assuming that the rate being regulated is fixed and constant (as described in Table I), but instead assuming that it is either a monotonically increasing or decreasing function of protein count. We perform this procedure for each feedback architecture, as depicted in Fig. 1. In order to obtain analytical expressions for noise, we use the well-known *linear noise approximation*. This involves linearizing the rate about the steady-state average protein count 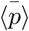 [54]. Note, these assumptions are only valid for small fluctuations in protein counts. This holds as regulatory protein levels in cells are tightly maintained within certain bounds. After linearizing the protein dependant rate, this new term is then substituted into the differential equations to replace the original constant rate that was associated with the gene-expression model without regulation in (2).

### A. Protein-Mediated Transcriptional Bursting Inhibition

To implement the first form of feedback, transcriptional bursting regulation (feedback architecture I in Fig. 1), we assume that transcriptional bursting events occur as a monotonically decreasing function *k*_*r*_(*p*(*t*)) of the protein count *p*(*t*). This results in a negative feedback strategy where an increase (decrease) in protein count will result in a decrease (increase) in bursting. Following the convention in [24], the rate *k*_*r*_(*p*(*t*)) is linearized about the steady-state average number of protein 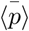 and we assume

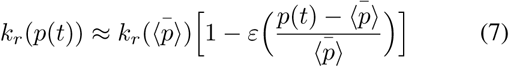

with an average transcription rate of 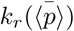. The dimension-less constant:

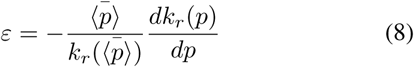

corresponds to the sensitivity of the transcription rate to changes in *p*(*t*) [24]. In simpler terms, this represents the strength of the feedback.

### B. Protein-Mediated Enhancement of pre-mRNA Degradation

Next, regulation by protein enhancement of pre-mRNA degradation is incorporated into the model (feedback architecture II in Fig. 1). In this feedback strategy, pre-mRNA degrades at a rate of *γ*_*r*_(*p*(*t*)). This form of regulation is more complex because it is not only dependent on protein count *p*(*t*), but also on pre-mRNA count *r*(*t*). We model this by assuming *γ*_*r*_(*p*(*t*))*r*(*t*) is a monotonically increasing function of protein count *p*(*t*). The approximated rate is linearized around both the average pre-mRNA count 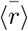 and protein count 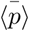. Again, it is assumed that *r*(*t*) and *p*(*t*) are tightly regulated and have small fluctuations from their respective steady-state averages. We find

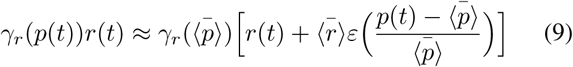

with an average pre-mRNA degradation rate 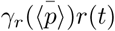 and dimensionless constant:

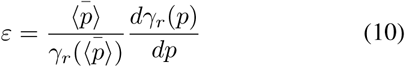

that is the sensitivity of the degradation rate to changes in *p*(*t*).

### C. Protein-Mediated Inhibition of pre-mRNA Export

Last, we incorporate the regulation of pre-mRNA export from the cell nucleus into the cytoplasm (feedback architecture III in Fig. 1). Here, we assume that pre-mRNA export is a monotonically decreasing function based on protein count, but is also dependant on pre-mRNA count. Thus, the total export rate *k*_*m*_(*p*(*t*))*r*(*t*) depends on both *p*(*t*) and *r*(*t*). The linear approximation about 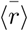 and 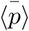 assumes that *r*(*t*) and *p*(*t*) are tightly regulated and have small fluctuations from their steady-state averages. We approximate the total export rate as:

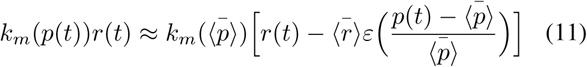

with an average export rate of 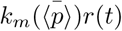 and dimension-less constant:

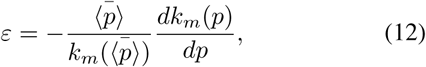

that is the sensitivity of the export rate to changes in *p*(*t*).

The time evolution of the statistical moments of each feedback mechanism are obtained by substituting the corresponding average rate found by linear approximation for the original rate in (1). This results in a unique solution of moment dynamics for each feedback architecture I-III. Based on the moment dynamics, the resulting protein noise levels of each feedback architecture are quantified as 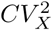, *Xϵ* {I, II, III} and displayed in the middle column of Fig. 2 (coefficient of variation squared of architecture *X*). Additionally, when there is no feedback strength, *ε* = 0, the resulting noise corresponds to protein level noise when there is no feedback present 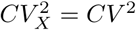.

***Remark 1:*** A close examination of the gene-expression noise equations in Fig. 2 reveals that noise can become unbounded for some values of feedback strength (*ε*). Specifically, it can be seen that 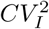 and 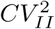 will remain stable as long as the feedback strength (*ε*) is satisfied:

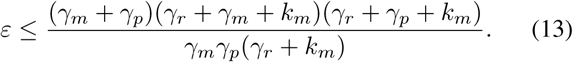

In contrast, 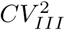 remains stable for all values of *ε >* 0.

## IV. COMPARISON BETWEEN FEEDBACK STRATEGIES

Mathematically controlled comparisons are performed to determine which feedback architecture best attenuates noise in gene-expression. We control these comparisons by assuming the same steady-state average values among the three feedback strategies. An obvious, yet important, trend is that noise is always reduced when regulation is incorporated. Any feedback strategy, in any parameter regime, outperforms the gene-expression model without feedback.

The first observation is made by considering a highly unstable pre-mRNA, thus pre-mRNA with a high degradation rate. When comparing regulation of transcription to regulation of pre-mRNA degradation, we find:

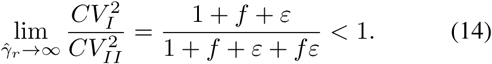

This ratio reveals that for a fixed feedback strength (*ε*), feedback architecture I outperforms feedback architecture II. However, when comparing regulation of pre-mRNA degradation to regulation of pre-mRNA export, the following result is found:

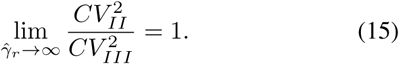

Under this condition, the noise reduction of feedback architectures II and III is the same, regardless of feedback strength. Based on these observations, we conclude that when pre-mRNA is unstable, feedback architecture I will always provide better noise attenuation than feedback architecture II and III, as well as the no feedback case. We can additionally analyze this regime in relation to feedback strength by selecting an increased pre-mRNA degradation rate and setting the remaining parameters equal to one. This is illustrated in Fig. 3D. We again conclude that transcriptional bursting feedback (feedback architecture I) is best.

**Fig. 3:**
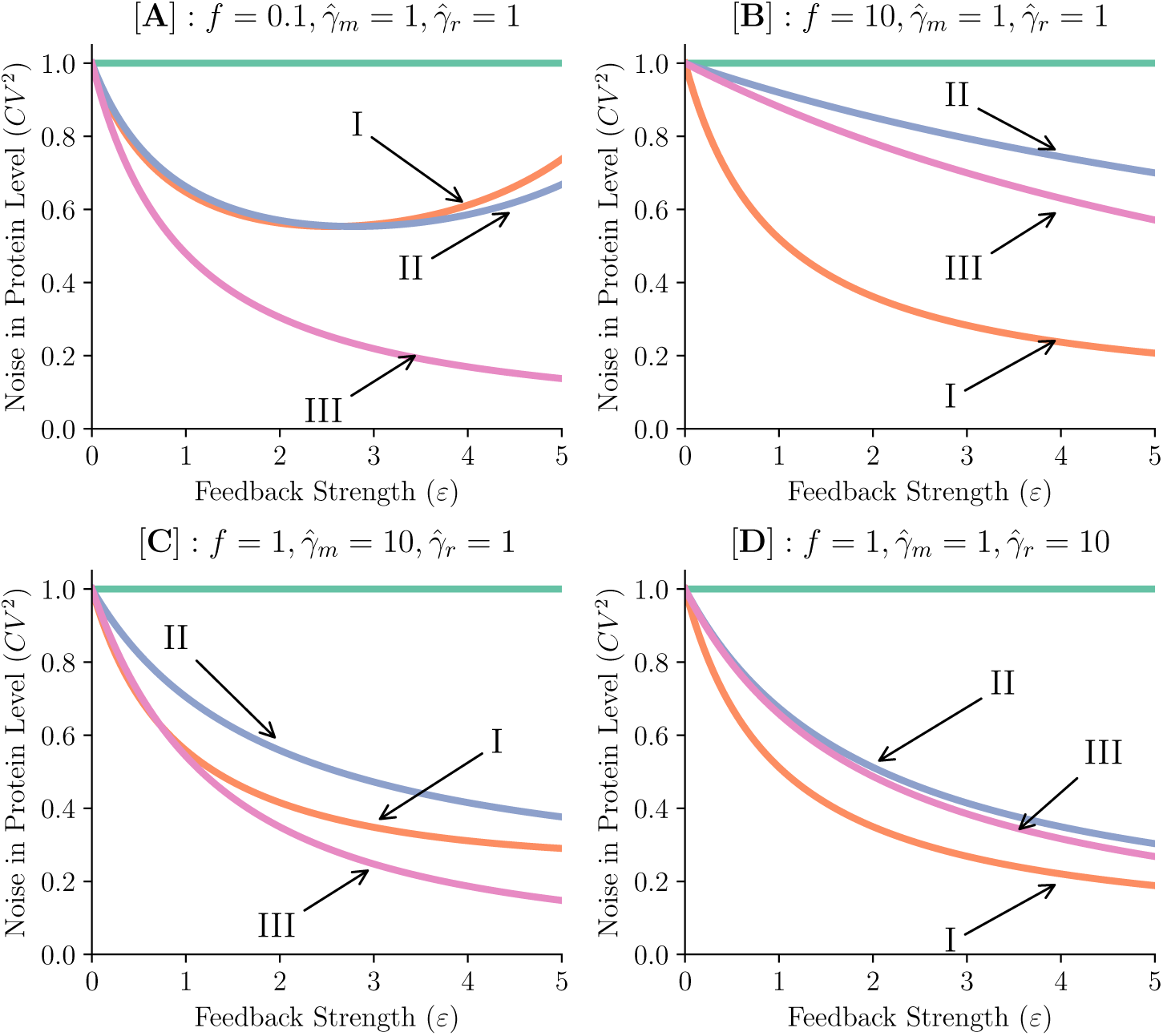
Comparison of noise in protein level for the three feedback strategies considered in the paper. Noise of each feedback architecture, normalized by noise of the no feedback case (green), is plotted as a function of increasing feedback strength (*ϵ*). Four cases of interest corresponding to different values of *f*, 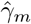, and 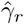 are shown while the remaining parameters are held constant: ⟨*B*_*e*_⟩ = 10 and 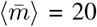. [A]: When *f <* 1, representing the case of pre-mRNA to more likely decay than be exported from the nucleus, feedback architecture III is best. [B]: When *f >* 1, feedback architecture I provides the most noise attenuation. [C]: When mRNA is unstable 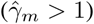, performance is dependant on feedback strength. At low strength, feedback architectures I and III behave similarly, but as feedback strength increases, feedback architecture III outperforms I and II. [D]: When pre-mRNA is unstable 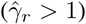, feedback architecture I suppresses noise the most.

Next, we compare noise suppression of each feedback architecture in relation to pre-mRNA processing. In the nucleus, retained pre-mRNA will either degrade or be exported into the cytoplasm. When the pre-mRNA degradation rate is higher than its export rate, feedback architecture III provides the best noise suppression. On the contrary, when pre-mRNA export is much higher than its degradation, feedback architecture I provides far superior noise suppression compared to feedback architectures II and III. Examples of noise suppression’s dependence on *f* are illustrated in Fig. 3A and 3B.

We last consider the dependence of noise suppression on mRNA stability. We model an unstable mRNA by selecting a high mRNA degradation rate, and setting the remaining parameters equal to one (illustrated in Fig. 3C). Here, we find feedback of pre-mRNA export, feedback architecture III, to provide the most noise suppression.

## V. CONCLUSIONS

This paper explores possible regulatory motifs utilized by cells to reduce the inherent stochasticity of gene-expression. Explicit analytical equations were developed to compare noise suppression of three possible feedback architectures (Fig. 2). After investigation through mathematically controlled comparisons, it is discovered that the best feedback architecture is dependant on the various parameters associated with gene-expression (Fig. 3). Transcriptional feedback (feedback architecture I) provides the best noise regulation when pre-mRNA export occurs more than its degradation, or when pre-mRNA is unstable. Additionally, in regimes when pre-mRNA degradation is more likely than its export, or when mRNA is unstable, then feedback of pre-mRNA export (feedback architecture III) is best.

Insight gained from feedback implemented at the pre-mRNA level can be related to post-transcriptional control of gene-expression in HIV. Inside the nucleus, pre-mRNA undergoes splicing and generates both incompletely and fully spliced transcripts. The incompletely spliced transcripts are retained in the nucleus while the fully spliced transcripts are exported into the cell’s cytoplasm to encode the regulatory proteins Tat, Rev, and Nef [55]–[58]. Interestingly, the Rev protein has the ability to bind to the incompletely spliced transcripts to allow for their export out of the nucleus [59]. This Rev mediated export decreases retained pre-mRNAs, which results in less fully spliced mRNA. Ultimately, the production of downstream proteins, including Rev, is reduced. With time, this proceeds as a negative feedback loop that we can relate to feedback architecture II described in this work [60]–[62].

Biological systems include time delays that alter stability by creating realistic oscillations in the system. Incorporation of these intrinsic delays will allow further investigation of noise suppression properties under more physiologically relevant conditions. Future work will include the addition of time delays to the gene-expression model proposed in this paper and resulting analysis of feedback strategies I-III performance.

## ACKNOWLEDGMENT

This work was supported by National Science Foundation through grant ECCS–1711548 to AS.

## REFERENCES

[1] W. J. Blake, M. Kaern, C. R. Cantor, and J. J. Collins, “Noise in eukaryotic gene expression,” Nature, vol. 422, pp. 633–637, 2003.

[2] A. Bar-Even, J. Paulsson, N. Maheshri, M. Carmi, E. OtShea, Y. Pilpel, and N. Barkai, “Noise in protein expression scales with natural protein abundance,” Nature Genetics, vol. 38, pp. 636–643, 2006.

[3] P. Bokes, Y. Ting Lin, and A. Singh, “High cooperativity in negative feedback can amplify noisy gene expression,” Bulletin for Mathematical Biology, vol. 80, pp. 1871–1899, 2018.

[4] S. M. Shaffer, M. C. Dunagin, S. R. Torborg, E. A. Torre, B. Emert, C. Krepler, M. Beqiri, K. Sproesser, P. A. Brafford, M. Xiao, E. Eggan, I. N. Anastopoulos, C. A. Vargas-Garcia, A. Singh, K. L. Nathanson, M. Herlyn, and A. Raj, “Rare cell variability and drug-induced reprogramming as a mode of cancer drug resistance,” Nature, vol. 546, pp. 431–435, 2018.

[5] E. Libby, T. J. Perkins, and P. S. Swain, “Noisy information processing through transcriptional regulation,” Proceedings of the National Academy of Sciences, vol. 104, pp. 7151–7156, 2007.

[6] H. B. Fraser, A. E. Hirsh, G. Giaever, J. Kumm, and M. B. Eisen, “Noise minimization in eukaryotic gene expression,” PLOS Biology, vol. 2, p. e137, 2004.

[7] B. Lehner, “Selection to minimise noise in living systems and its implications for the evolution of gene expression,” Molecular Systems Biology, vol. 4, p. 170, 2008.

[8] R. Kemkemer, S. Schrank, W. Vogel, H. Gruler, and D. Kaufmann, “Increased noise as an effect of haploinsufficiency of the tumor-suppressor gene neurofibromatosis type 1 in vitro,” Proceedings of the National Academy of Sciences, vol. 99, pp. 13783–13788, 2002.

[9] D. L. Cook, A. N. Gerber, and S. J. Tapscott, “Modeling stochastic gene expression: implications for haploinsufficiency,” Proceedings of the National Academy of Sciences, vol. 95, pp. 15641–15646, 1998.

[10] R. Bahar, C. H. Hartmann, K. A. Rodriguez, A. D. Denny, R. A. Busuttil, M. E. Dolle, R. B. Calder, G. B. Chisholm, B. H. Pollock, C. A. Klein, and J. Vijg, “Increased cell-to-cell variation in gene expression in ageing mouse heart,” Nature, vol. 441, pp. 1011–1014, 2006.

[11] A. Brock, H. Chang, and S. Huang, “Non-genetic heterogeneity – a mutation-independent driving force for the somatic evolution of tumours,” Nature Reviews Genetics, vol. 10, pp. 336–342, 2009.

[12] N. Balaban, J. Merrin, R. Chait, L. Kowalik, and S. Leibler, “Bacterial persistence as a phenotypic switch,” Science, vol. 305, pp. 1622–1625, 2004.

[13] J. Feng, D. A. Kessler, E. Ben-Jacob, and H. Levine, “Growth feedback as a basis for persister bistability,” Proceedings of the National Academy of Sciences, vol. 111, pp. 544–549, 2014.

[14] I. E. Meouche, Y. Siu, and M. J. Dunlop, “Stochastic expression of a multiple antibiotic resistance activator confers transient resistance in single cells,” Scientific Reports, vol. 6, p. 19538, 2016.

[15] A. Singh and L. S. Weinberger, “Stochastic gene expression as a molecular switch for viral latency,” Current Opinion in Microbiology, vol. 12, pp. 460–466, 2009.

[16] M. Voliotis and C. G. Bowsher, “The magnitude and colour of noise in genetic negative feedback systems,” Nucleic Acids Research, 2012.

[17] A. Singh and J. P. Hespanha, “Optimal feedback strength for noise suppression in autoregulatory gene networks,” Biophysical Journal, vol. 96, pp. 4013–4023, 2009.

[18] H. El-Samad and M. Khammash, “Regulated degradation is a mechanism for suppressing stochastic fluctuations in gene regulatory networks,” Biophysical Journal, vol. 90, pp. 3749–3761, 2006.

[19] A. Singh and J. P. Hespanha, “Evolution of autoregulation in the presence of noise,” IET Systems Biology, vol. 3, pp. 368–378, 2009.

[20] I. Lestas, G. Vinnicombe, and J. Paulsson, “Fundamental limits on the suppression of molecular fluctuations,” Nature, vol. 467, pp. 174–178, 2010.

[21] P. S. Swain, “Efficient attenuation of stochasticity in gene expression through post-transcriptional control,” Journal of Molecular Biology, vol. 344, pp. 956–976, 2004.

[22] J. M. Pedraza and J. Paulsson, “Effects of molecular memory and bursting on fluctuations in gene expression,” Science, vol. 319, pp. 339–343, 2008.

[23] Y. Morishita and K. Aihara, “Noise-reduction through interaction in gene expression and biochemical reaction processes,” Journal of Theoretical Biology, vol. 228, pp. 315–325, 2004.

[24] A. Singh, “Negative feedback through mRNA provides the best control of gene-expression noise,” IEEE Transactions on Nanobioscience, vol. 10, pp. 194–200, 2011.

[25] A. Singh and J. P. Hespanha, “Reducing noise through translational control in an auto-regulatory gene network,” in Proc. of the 2009 Amer. Control Conference, St. Louis, MO, 2009.

[26] P. Bokes and A. Singh, “Gene expression noise is affected deferentially by feedback in burst frequency and burst size,” Journal of Mathematical Biology, vol. 74, pp. 1483–1509, 2017.

[27] A. Borri, P. Palumbo, and A. Singh, “The impact of negative feedback in metabolic noise propagation,” IET Systems Biology, pp. 179–186, 2016.

[28] Y. Dublanche, K. Michalodimitrakis, N. Kummerer, M. Foglierini, and L. Serrano, “Noise in transcription negative feedback loops: simulation and experimental analysis,” Molecular Systems Biology, vol. 2, p. 41, 2006.

[29] D. Nevozhay, R. M. Adams, K. F. Murphy, K. Josic, and G. Balazsi, “Negative autoregulation linearizes the dose response and suppresses the heterogeneity of gene expression,” Proceedings of the National Academy of Sciences, vol. 106, pp. 5123–5128, 2009.

[30] M. Pons, S. Prieto, L. Miguel, T. Frebourg, D. Campion, C. Suñé, and M. Lecourtois, “Identification of TCERG1 as a new genetic modulator of TDP-43 production in Drosophila,” Acta Neuropathologica Communications, vol. 6, 2018.

[31] A. Singh and P. Bokes, “Consequences of mRNA transport on stochastic variability in protein levels,” Biophysical Journal, vol. 103, pp. 1087–1096, 2012.

[32] P. Konieczny, E. Stepniak-Konieczna, and K. Sobczak, “MBNL expression in autoregulatory feedback loops,” RNA Biology, vol. 15, pp. 1–8, 2018.

[33] M. Sturrock, S. Li, and V. Shahrezaei, “The influence of nuclear compartmentalisation on stochastic dynamics of self-repressing gene expression,” Journal of Theoretical Biology, vol. 424, pp. 55–72, 2017.

[34] M. Aprile, S. Cataldi, M. R. Ambrosio, V. D’Esposito, K. Lim, A. Dietrich, M. Bluher, D. B. Savage, P. Formisano, A. Ciccodicola, and V. Costa, “PPAR*γd*5, a naturally occurring dominant-negative splice isoform, impairs PPAR*γ* function and adipocyte differentiation,” Cell, vol. 25, pp. 1577–1592, 2018.

[35] M. M. Hansen, W. Y. Wen, E. Ingerman, B. S. Razooky, C. E. Thompson, R. D. Dar, C. W. Chin, M. L. Simpson, and L. S. Weinberger, “A post-transcriptional feedback mechanism for noise suppression and fate stabilization,” Cell, vol. 173, pp. 1609–1621, 2018.

[36] T. Stoeger, N. Battich, and L. Pelkmans, “Passive noise filtering by cellular compartmentalization,” Cell, vol. 164, pp. 1151–1161, 2016.

[37] N. Battich, T. Stoeger, and L. Pelkmans, “Control of transcript variability in single mammalian cells,” Cell, vol. 163, pp. 1596–1610, 2015.

[38] M. M. Hansen, R. V. Desai, M. L. Simpson, and L. S. Weinberger, “Cytoplasmic amplification of transcriptional noise generates substantial cell-to-cell variability,” Cell, vol. 7, pp. 384–397, 2018.

[39] N. Kumar, A. Singh, and R. V. Kulkarni, “Transcriptional bursting in gene expression: analytical results for general stochastic models,” PLoS Computational Biology, vol. 11, p. e1004292, 2015.

[40] A. Singh, B. Razooky, C. D. Cox, M. L. Simpson, and L. S. Weinberger, “Transcriptional bursting from the HIV-1 promoter is a significant source of stochastic noise in HIV-1 gene expression,” Biophysical Journal, vol. 98, pp. L32–L34, 2010.

[41] R. D. Dar, B. S. Razooky, A. Singh, T. V. Trimeloni, J. M. McCollum, C. D. Cox, M. L. Simpson, and L. S. Weinberger, “Transcriptional burst frequency and burst size are equally modulated across the human genome,” Proceedings of the National Academy of Sciences, vol. 109, pp. 17454–17459, 2012.

[42] A. Raj, C. S. Peskin, D. Tranchina, D. Vargas, and S. Tyagi, “Stochastic mRNA synthesis in mammalian cells,” PLOS Biology, vol. 4, p. e309, 2006.

[43] I. Golding, J. Paulsson, S. Zawilski, and E. Cox, “Real-time kinetics of gene activity in individual bacteria,” Cell, vol. 123, pp. 1025–1036, 2005.

[44] D. M. Suter, N. Molina, D. Gatfield, K. Schneider, U. Schibler, and F. Naef, “Mammalian genes are transcribed with widely different bursting kinetics,” Science, vol. 332, pp. 472–474, 2011.

[45] S. Yunger, L. Rosenfeld, Y. Garini, and Y. Shav-Tal, “Single-allele analysis of transcription kinetics in living mammalian cells,” Nature Methods, vol. 7, pp. 631–633, 2010.

[46] C. Fritzsch, S. Baumgärtner, M. Kuban, D. Steinshorn, G. Reid, and S. Legewie, “Estrogen-dependent control and cell-to-cell variability of transcriptional bursting,” Molecular Systems Biology, vol. 14, p. e7678, 2018.

[47] M. A. Savageau, Biochemical Systems Analysis: A Study of Function and Design in Molecular Biology. Addison-Wesley, Reading, MA, 1976.

[48] J. P. Hespanha and A. Singh, “Stochastic models for chemically reacting systems using polynomial stochastic hybrid systems,” International Journal of Robust and Nonlinear Control, vol. 15, pp. 669–689, 2005.

[49] A. Singh and J. P. Hespanha, “Approximate moment dynamics for chemically reacting systems,” IEEE Transactions on Automatic Control, vol. 56, pp. 414–418, 2011.

[50] J. Paulsson, “Summing up the noise in gene networks,” Nature, vol. 427, pp. 415–418, 2004.

[51] A. Becskei and L. Serrano, “Engineering stability in gene networks by autoregulation,” Nature, vol. 405, pp. 590–593, 2000.

[52] D. Bergeron, G. Pal, Y. B. Beaulieu, B. Chabot, and F. Bachand, “Regulated intron retention and nuclear pre-mRNA decay contribute to PABPN1 autoregulation,” Molecular and Cellular Biology, vol. 35, pp. 2503–2517, 2015.

[53] R. Bastos, A. Lin, M. Enarson, and B. Burke, “Targeting and function in mRNA export of nuclear pore complex protein Nup153,” Journal of Cell Biology, vol. 134, pp. 1141–1156, 1996.

[54] N. Van Kampen, Stochastic processes in physics and chemistry. Elsevier, 2011.

[55] S. Kim, R. Byrn, J. Groopman, and D. Baltimore, “Temporal aspects of DNA and RNA synthesis during human immunodeficiency virus infection: evidence for differential gene expression,” *Journal of Virol-* infection: evidence for differential gene expression,” Journal of Virology, vol. 63, pp. 3708–3713, 1989.

[56] D. Purcell and M. Martin, “Alternative splicing of human immunodeficiency virus type 1 mRNA modulates viral protein expression, replication, and infectivity,” Journal of Virology, vol. 67, pp. 6365–6378, 1993.

[57] B. Rojas-Araya, T. Ohlmann, and R. Soto-Rifo, “Translational control of the HIV unspliced genomic RNA,” Viruses, vol. 7, pp. 4326–4351, 2015.

[58] R. Pomerantz, D. Trono, M. Feinberg, and D. Baltimore, “Cells nonproductively infected with HIV-1 exhibit an aberrant pattern of viral RNA expression: a molecular model for latency,” Cell, vol. 61, pp. 1271–1276, 1990.

[59] M. Malim, J. Hauber, S. Le, J. Maizel, and B. Cullen, “The HIV-1 Rev trans-activator acts through a structured target sequence to activate nuclear export of unspliced viral mRNA,” Nature, vol. 338, pp. 254–257, 1989.

[60] K. Kim and J. Yin, “Robust growth of human immunodeficiency virus type 1 HIV-1,” Biophysical Journal, vol. 89, pp. 2210–2221, 2005.

[61] M. Malim, J. Hauber, R. Fenrick, and B. Cullen, “Immunodeficiency virus Rev trans-activator modulates the expression of the viral regulatory genes,” Nature, vol. 335, pp. 181–183, 1988.

[62] B. K. Felber, C. M. Drysdale, and G. N. Pavlakis, “Feedback regulation of human immunodeficiency virus type 1 expression by the Rev protein,” Journal of Virology, vol. 64, p. 3734, 1990.

